# Skin aging risk factors: A nationwide population study in Mongolia Risk factors of skin aging

**DOI:** 10.1101/2021.03.22.436407

**Authors:** Tuya Nanzadsuren, Tuul Myatav, Amgalanbaatar Dorjkhuu, Mandukhai Ganbat, Chuluunbileg Batbold, Baljinnyam Batsuuri, Khandsuren Byamba

## Abstract

The world population is aging and no country is immune to the consequences. We are not aware of any country-specific skin aging risk factors data for the Mongolian people. Thus, we aimed to study the risk factors associated with skin aging in the Mongolian population. A population-based cross-sectional study of 2720 study participants 18 years of age and older was performed evaluating the severity of skin aging based on cutaneous microtopography. Questionnaire data and skin physiological measurements were obtained. The odds ratios for skin aging grades associated with risk factors were estimated using ordinal logistic regression. Study participant’s mean age was 45 years, ranging from 18 to 87. After adjustment for known risk factors, skin aging was associated with demographic risk factors such as increasing age (aOR=1.19, 95% CI 1.18-1.20), living in an urban area (aOR=1.31, 95% CI 1.12-1.55) and working outside (aOR=1.44, 95% CI 0.88-2.39) and lifestyle factors including non-usage of sunscreen cream (aOR=1.09 95% CI 0.87-1.37), being a smoker (aOR=1.32, 95% CI 1.09-1.61), having a higher body mass index (BMI) (aOR=1.04, 95% CI 1.02-1.06) and higher levels of sun exposure time (aOR=1.03, 95% CI 1.00-1.06) were significantly associated with higher skin aging grades. Having dry (aOR=1.94, 95% CI 1.45-2.59) and combination skin (aOR=1.62, 95% CI 1.22-2.16) types were also independent risk factors associated with skin aging. Having very low skin surface moisture at the T-zone (aOR=2.10, 95% CI 1.42-3.11) and U-zone (aOR=1.25, 95% CI 0.95-1.65) were significantly related to skin aging. Older age, urban living, harsh working conditions, living in a ger district were independent demographic risk factors related to skin aging. Not using sunscreen cream, smoking, higher BMI, greater levels of sun exposure were significant lifestyle risk factors. Having a skin type other than normal was a physiologic risk factor for skin aging.

## Introduction

The aging of the worldwide population is now widely recognized as an issue that impacts skin [1]. Skin aging is affected by demographic, environmental, and lifestyle factors and skin types [2].

Demographic factors play an important role in skin aging. In 2019, a Nepal study with Caucasian or Mongolian tribes demonstrated that increasing age and sun exposure were the main determinants of skin aging [3]. Sun exposure on the visible areas of the skin affects the progression of skin aging by up to 80% [4]. A study demonstrated that men had significantly higher skin aging grades than women in all age groups [5]. A study of 1204 Indian women living in Chennai city, the fourth most populous urban area in India, found that living in a metropolitan area was a risk factor of skin aging [6]. Skin is vulnerable to work conditions, and outdoor work, even part-time, leads to skin aging [5, 7].

Lifestyle factors are primarily preventable causes of skin aging, which is the central theme for skin health. In 2014, an American study identified that poor quality sleepers had more advanced skin aging signs than good sleepers [8]. Increasing pack-years of smoking significantly impacted skin wrinkles and skin aging, observed in a worldwide internet-based study [9]. Some studies observed that sunscreen and wrinkle prevention products positively affected skin aging [10, 11]. Menopausal and postmenopausal women were significantly more likely to have higher skin aging grades than premenopausal women [12]. Skin type is also a significant factor in skin aging. People who have dry skin are more vulnerable to having wrinkles than oily skin [13]. Furthermore, some skin biomarkers, such as melatonin and epidermal growth factor, are associated with fewer wrinkles and less aging [14, 15].

Most studies of factors relating to skin aging have focused on the developed world, and few have examined the risk factors in developing nations. It is unclear how Mongolia’s long cold winter with little sunlight, extensive urban air pollution, and its peoples’ diet of primarily meat and milk impacts their skin aging. Our study aimed to investigate the relationship between skin aging and its potential risk factors using non-invasive cutaneous microtopography in what we believe is Mongolia’s first nation-wide skinaging survey.

## Materials and methods

### Study design

We employed a nationwide cross-sectional study design.

### Study subjects and sampling

Our study was conducted in 2018 between May and August. A total of 2731 people, 18 years and older, agreed to participate in this study, with 2720 completing the survey. The response rate was 99.6%. The study population was relatively healthy adults randomly selected from Western, Eastern, Northern, and Central regions of Mongolia and Mongolia’s capital, Ulaanbaatar. This geographically diverse sample represented the ethnic and socioeconomic characteristics of the Mongolian national population. The study exclusion criteria were having a history of skin surgery, dermatitis and skin infectious disease.

### Data collection method

#### Questionnaire

A questionnaire was developed as part of our study. We established a committee of experts from the Department of Dermatology and biostatistics team from the School of Medicine, Mongolian National University of Medical Sciences. Several items from an existing questionnaire from the Department of Preventive Medicine, School of Public Health were used. Additionally, we developed new questions regarding lifestyle factors with two to five responses and determined questionnaire length. We verified the questionnaire item’s face validity in terms of readability and understandability by the study participants with a pilot study of twenty participants. The 41-item questionnaire took approximately fifteen minutes to complete. The questionnaire in both Mongolian and English language is attached as S1 appendix.

Each participant was asked to complete the questionnaire to gather some personal socioeconomic data, and lifestyle and behavioral risk factors potentially contributing to aging skin. Information about personal characteristics included: age, gender, education level, living area, type of housing, BMI, dietary habits, sleep time per night, smoking status, work conditions, tobacco consumption, sunscreen usage, frequency of getting professional skin care services and menopausal status.

#### Skin physiological parameters

We obtained the skin physiological parameters using the Skin Diagnosis System SD-Pro (BOMTECH Co., LTD, Korea) [16]. This device incorporates a moisture checker, sebum tape, digital camera with a light meter and microscope. We used the moisture checker and sebum tape to quantify skin surface moisture and sebum at the T-zone (T-shaped area of the forehead, nose and chin) and U-zone (U-shaped area of the temple and cheeks). The skin in these zones differs in that T-zone has more sebaceous glands than the U-zone. The device measures the skin surface moisture in units of percent, ranging from 1.0% to 45.0%. The skin surface sebum was measured by the number of oil content in per unit area. The width of eye wrinkles and nose pores were measured with digital calipers on the device using standardized digital photographs of the participants’ eyes and nose. Using cheek and chin photographs, the skin pigmentation was classified into one of 3 grades by comparison to nine standard images on the machine.

#### Blood samples

A total of 5 ml of blood was collected via capillary puncture from each participant between 8 am and 9 am. The blood sample was centrifuged at room temperature at 1500 revolutions per minute for 10 minutes. The melatonin and epidermal growth factor (EGF) levels were measured using a semi-automated enzyme-linked immunosorbent assay analyzer (Dynex Technologies DS2^®^; DRG International, Germany) [17]. Serum melatonin and EGF quantitative measurements were expressed as ng/L and pg/mL.

#### Skin aging grade

We used cutaneous microtopography to evaluate skin aging grades. The subject lightly clenched their left hand and placed it on an incline board at a 30° angle of inclination. Digital photographs were then taken of the participant’s dorsal hand area using a standardized process. We determined the skin aging grades by comparing our photographs to the standard images in the Beagley-Gibson system [18, 19]. This six-step grading system is based on progressive alterations in the skin surface characteristics associated with actinic changes in standard photographs. Representative dorsal hand images of nine study participants classified using the Beagley-Gibson method are shown in Figure 1. Grade 1 is characterized by primary lines that were all the same depth. Secondary lines were clearly visible, nearly the same depth as the primaries, and often met to form an apex of triangles (“star formation”). Grade 2 has some flattening and loss of clarity of the secondary lines. Star formations were still present, but often one or more of the secondary lines that form the configuration is unclear. Grade 3 is characterized by unevenness of the primary lines and noticeable flattening of the secondary lines with little or no star formation. Grade 4 has macroscopic deterioration in texture and coarse, deep primary lines, with distortion and loss of secondary lines. Grade 5 has noticeable flat skin between the primary lines, with few or no secondary lines. Grade 6 has large, deep, and widely spaced primary lines, with no secondary lines.

**Fig 1.**
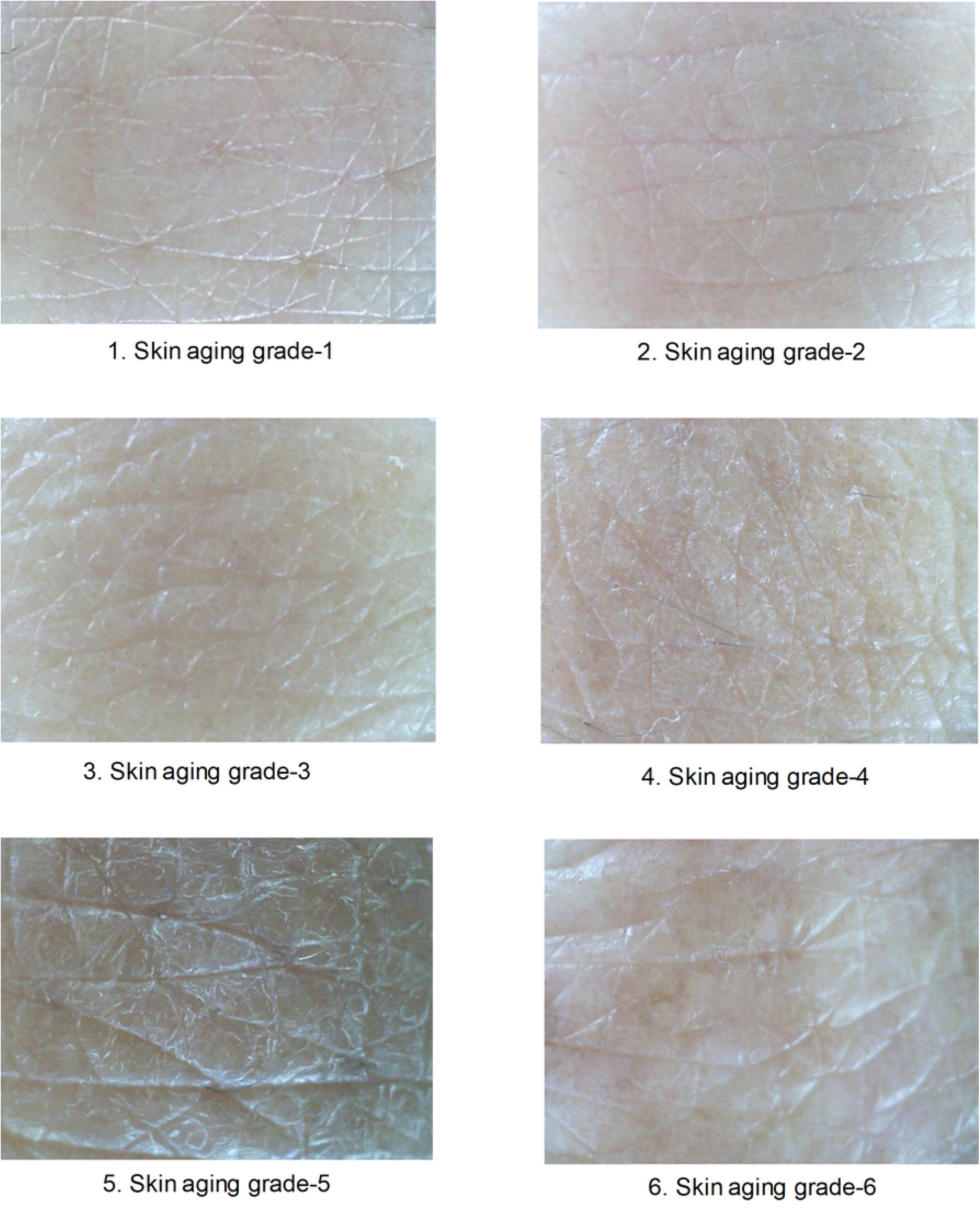
Representative dorsal hand images of nine study participants classified using the Beagley-Gibson method.

#### Ethical considerations

The study was approved by the Mongolian National University of Medical Sciences and the biomedical ethical committee on April 25, 2013 (No. 13-12/1A). Written informed consent was obtained from each participant.

#### Statistical methods

The dependent variable in this study was skin aging which is ordinal data scaled from 2 to 6 skin-aging grades according to Beagley Gibson method. Gender, work conditions, smoking status, sunscreen usage, geographic location, housing type, frequency of getting professional skin care services, skin type, night sleep time and menstrual status were independent categorical variables. Additionally, the participant’s age, BMI, time of sun exposure, and serum melatonin and epidermal growth factor levels were continuous data. We determined potential skin physiological risk factors associated with skin aging in skin surface moisture and sebum at T- and U-zones, skin pore size, pigmentation, and wrinkles, which were all continuous data. We presented continuous variables as the mean and standard deviation and determined that they were normally distributed using the Shapiro-Wilk test. Categorical data were summarized as frequencies and percentages.

Spearman’s correlation analysis was used to determine the correlation between skin aging grades and the skin physiological parameters, including skin surface moisture and sebum at the T- and U-zone, skin pore size and wrinkle, age, BMI, sun exposure time, serum melatonin and epidermal growth factor levels.

To assess the association between skin aging grades and potential risk factors contributing to skin aging, odds ratios (OR) and 95% confidence intervals (CI) were estimated using ordinal logistic regression models with proportional odds assumption. Our ordinal logistic regression model assumed that as the independent variables increased, the probability of falling into the next higher skin aging grade increased. The odds ratios are interpreted depending on the type of variable. For continuous variables, the variables predict the odds of being in the next higher skin-aging grade per unit increase in the predictor variable while holding the other predictor variables constant. For nominal predictor variables, the estimated odds ratio predicts the odds of being in a given skin aging grade compared to the reference category. We calculated both crude odds ratio and adjusted odds ratio. The crude odds ratio expressed the independent variable’s odd ratio for predicting the dependent variable using an odds ratio calculation. The adjusted odds ratio was derived from our final ordinal regression model. The predictor variables were tested to verify the multicollinearity assumption was tenable. All statistical tests were 2-sided, with a p-value<0.05 considered significant. Stata 14 was used for carrying out all statistical analyses.

## Results

### Demographic factors

A total of 2731 people, over 18 years, were invited to take part in the study. Of those who agreed to participate in the study, 2720 completed the survey, as shown in Table 1. We determined serum melatonin and epidermal growth factor levels in 541 study participants. The mean age of the participants was 45, ranging from 18 to 87 years. Over 30% of the study participants had grade 2 and 3 skin aging, and almost 70% had grade 4 to 6 skin aging. Women were 54% of the sample. Over 75% of the study participants worked in optimal conditions. Grade 6 skin aging was found in 27.6% of people living in urban and 14.6% of people living in rural areas.

**Table 1.** A Correlation between demographic and life style factors and skin aging grades.

| Variables | Skin aging grade |  |  |  |  | cOR | 95% CI | aOR* | 95% CI |
| --- | --- | --- | --- | --- | --- | --- | --- | --- | --- |
|  | 2 N (%) | 3 N (%) | 4 N (%) | 5 N (%) | 6 N (%) |  |  |  |  |
| Age, years<sup>¶</sup> | 29.3±10 | 32.8±8 | 41.4±8 | 52.2±9 | 61.8±8 | 1.03 | 1.01-1.05 | 1.19 | 1.18-1.20 |
| **Gender** |  |  |  |  |  |  |  |  |  |
| Male | 162 (12.9) | 275 (21.9) | 273 (21.8) | 305 (24.4) | 237 (18.9) | 0.984 | 0.86-1.12 | 0.899 | 0.73-1.10 |
| Female | 233 (15.9) | 271 (18.5) | 294 (20.0) | 385 (26.2) | 285 (19.4) | 1 |  | 1 |  |
| **Work condition** |  |  |  |  |  |  |  |  |  |
| Nightshift | 3 (5.6) | 18 (33.3) | 15 (27.8) | 9 (16.7) | 9 (16.7) | 0.876 | 0.55-1.38 | 1.317 | 0.80-2.15 |
| Outside | 5 (8.9) | 5 (8.9) | 19 (33.9) | 18 (32.1) | 9 (16.1) | 1.302 | 0.83-2.03 | 1.448 | 0.88-2.39 |
| Heavy | 63 (14.9) | 87 (20.6) | 92 (21.8) | 111 (26.3) | 69 (16.4) | 0.93 | 0.7-1.12 | 1.117 | 0.90-1.38 |
| Toxic | 12 (9.8) | 17 (13.9) | 26 (21.3) | 39 (32.0) | 28 (23.0) | 1.48 | 1.07-2.04 | 1.469 | 1.03-2.09 |
| Optimal | 312 (15.1) | 419 (20.3) | 415 (20.1) | 513 (24.8) | 407 (19.7) | 1 |  | 1. |  |
| **Geographical area** |  |  |  |  |  |  |  |  |  |
| Urban | 126 (13.2) | 155 (16.2) | 181 (19.0) | 229 (24.0) | 264 (27.6) | 1.615 | 1.40-1.86 | 1.314 | 1.12-1.55 |
| Rural | 269 (15.2) | 391 (22.2) | 386 (21.9) | 461 (26.1) | 258 (14.6) | 1 |  | 1 |  |
| **Housing type** |  |  |  |  |  |  |  |  |  |
| Ger (circular felt-covered tent) | 108 (12.5) | 151 (17.5) | 188 (21.8) | 262 (30.4) | 152 (17.67) | 1.131 | 0.95-1.34 | 1.215 | 0.98-1.5 |
| Ger district house | 139 (15.5) | 171 (19.1) | 185 (20.7) | 222 (24.8) | 178 (19.9) | 1.017 | 0.86-1.20 | 1.082 | 0.89-1.32 |
| Dormitory | 25 (20.3) | 40 (32.5) | 27 (22.0) | 22 (17.9) | 9 (7.3) | 0.501 | 0.36-0.69 | 1.081 | 0.75-1.56 |
| Apartment | 121 (14.6) | 182 (22.0) | 165 (20.0) | 182 (22.0) | 176 (21.3) | 1 |  | 1 |  |
| **Tobacco consumption** |  |  |  |  |  |  |  |  |  |
| Yes | 91 (12.4) | 161 (21.9) | 152 (20.7) | 193 (26.3) | 137 (18.7) | 1.035 | .89-1.20 | 1.326 | 1.09-1.61 |
| No | 304 (15.3) | 385 (19.4) | 414 (20.9) | 496 (25.0) | 385 (19.4) | 1 |  | 1 |  |
| **Amount of sleep** |  |  |  |  |  |  |  |  |  |
| 4-5 hours | 29 (8.7) | 31 (9.3) | 43 (12.9) | 124 (37.1) | 107 (32.0) | 2.68 | 2.15-3.35 | 0.43 | 0.71-1.18 |
| 6-7 hours | 188 (14.5) | 273 (21.1) | 301 (23.3) | 306 (23.7) | 225 (17.4) | 1.05 | 0.42-1.22 | 0.84 | 0.72-1.01 |
| 8-10 hours | 178 (16.3) | 242 (22.1) | 223 (20.4) | 260 (23.8) | 190 (17.4) | 1 |  | 1 |  |
| **Skin type** |  |  |  |  |  |  |  |  |  |
| Combination skin | 164 (15.5) | 243(22.9) | 215 (20.3) | 248 (23.4) | 189 (17.8) | 1.266 | 0.99-1.62 | 1.620 | 1.22-2.16 |
| Oily | 49 (11.8) | 78 (18.7) | 102 (24.5) | 117 (28.1) | 71 (17.0) | 1.524 | 1.15-2.02 | 1.544 | 1.11-2.14 |
| Dry | 141 (13.9) | 173 (17.1) | 188 (18.5) | 279 (27.5) | 233 (23.0) | 1.754 | 1.36-2.25 | 1.940 | 1.45-2.59 |
| Normal | 41 (17.8) | 52 (22.6) | 62 (27.0) | 46 (20.0) | 29 (12.6) | 1 |  | 1 |  |
| **Sunscreen usage** |  |  |  |  |  |  |  |  |  |
| Never | 225 (13.2) | 319 (18.7) | 336 (19.7) | 460 (27.0) | 366 (21.5) | 1.427 | 1.18-1.72 | 1.095 | 0.87-1.37 |
| Often | 89 (15.9) | 124 (22.2) | 138 (24.7) | 131 (23.5) | 76 (13.6) | 0.978 | 0.79-1.22 | 1.304 | 1.02-1.66 |
| Always | 81 (17.8) | 103 (22.7) | 92 (20.3) | 98 (21.6) | 80 (17.6) | 1 |  | 1 |  |
| **Frequency of getting professional skin care services** |  |  |  |  |  |  |  |  |  |
| Never | 270 (13.0) | 380 (18.4) | 414 (20.0) | 548 (26.5) | 457 (22.1) | 3.166 | 2.12-4.69 | 2.071 | 1.30-3.30 |
| Rarely | 97 (17.1) | 146 (25.8) | 138 (24.4) | 124 (21.9) | 61 (10.8) | 1.792 | 1.19-2.70 | 2.186 | 1.35-3.53 |
| Regularly | 28 (32.9) | 20 (23.5) | 15 (17.6) | 18 (21.2) | 4 (4.7) | 1 |  | 1 |  |
| **Menopausal status** |  |  |  |  |  |  |  |  |  |
| Postmenopausal | 19 (5.5) | 8 (2.3) | 38 (11.0) | 151 (43.6) | 130 (37.6) | 18.78 | 13.6-25.8 | 0.725 | 0.47-1.18 |
| Perimenopausal | 23 (18.3) | 28 (22.2) | 34 (27.0) | 33 (26.2) | 8 (6.3) | 1.86 | 1.30-2.65 | 1.121 | 0.78-1.61 |
| Premenopausal | 118 (24.0) | 155 (31.6) | 132 (26.9) | 78 (15.9) | 8 (1.6) | 1 |  | 1 |  |

%mean±standard deviation, \*- adjusted for all variables in the model except from data of menopausal status

Table 1 also shows the odds ratios of the skin-aging grades. Age had the most impact on skin aging. On the adjusted analysis, increasing age by one year was associated with a 19% increase in odds of having a higher skin aging grade. There was no effect of gender on the skin-aging grade. Compared to Mongolians in rural areas, those in urban areas had almost 60% increased odds of higher grades (OR=1.61, 95% CI 1.40-1.86) on crude analysis. People working in harsh conditions were approximately 1.5 times more likely to have higher skin aging grades than others working in optimal conditions.

### Lifestyle factors

People with a normal skin type had a higher prevalence of grade 2 skin aging than people having dry, oily and combination skin types (p<0.001). The frequency of getting professional skin care service was strongly associated with skin aging grade. Those who never and rarely get professional skin care services had twice the odds (aOR=2.07, 95% CI 1.30-3.30, aOR=2.19, 95% CI 1.35-3.53) of having higher skin aging grades compared with those who often get professional skin care services. Similarly, after adjusting for all other variables, people who reported rarely and never using sunscreen cream had significantly elevated odds of higher skin aging grades (OR=1.30, 95% CI 1.02-1.66) compared with those who reported always using sunscreen cream. The odds of smokers having higher skin aging grades than non-smokers were 1.31, with a 95% CI of 1.09-1.61. There was no significant association between housing type and skin aging grade. Compared with those with a normal skin type, those with dry, oily and combination skin types had significantly increased odds of having higher skin aging grades (aOR=1.94 95% CI 1.45-2.59, aOR=1.54 95% CI 1.11-2.14 and aOR=1.62, 95% CI 1.22-2.16 respectively). On crude analysis, sleeping fewer hours was associated with an almost twofold risk of higher skin aging compared to 8 to 10 hours of night sleep. But this was not significant after adjusting all the variables. Increasing the time outside by 1 hour per day was associated with a nearly 3% increase in odds of a higher skin aging grade (aOR=1.03, 95% CI 1.00-1.06). Similarly, a 1 unit increase in the BMI increased the odds of being in the next skin aging category by 4% (aOR=1.04, 95% CI 1.02-1.06).

In our study, 963 of 1474 women completed the questions about their menstrual cycle. For this reason, we analyzed this variable using a separate model and did not include this menstrual status in our overall ordinal regression model. Postmenopausal and perimenopausal women had higher odds of skin aging than those with a regular menstrual cycle (aOR=1.12, 95% CI 0.78-1.61) (Table 1).

### Skin type

Table 2 shows skin physiological parameters, including skin surface sebum, moisture, pigmentation, wrinkles and skin pore size. Very low skin surface sebum was found in 27% of patients at the T-zone and 46% at the U-zone in those with grade 6 skin aging. Across all skin aging grades, 30-50% of study participants had medium levels of surface moisture at T- and U-zones. Approximately 70% of people with grade 6 skin aging had relatively large skin pores.

**Table 2.** An association between skin physiological measurements and skin aging grades.

| Variables | Skin aging grade N (%) |  |  |  |  | OR | 95%CI | aOR* | 95%CI |
| --- | --- | --- | --- | --- | --- | --- | --- | --- | --- |
|  | 2 | 3 | 4 | 5 | 6 |  |  |  |  |
| Skin surface sebum T-zone |  |  |  |  |  |  |  |  |  |
| Very low | 25 (6.3) | 20 (3.7) | 36 (6.4) | 60 (8.7) | 84 (16.1) | 2.50 | 1.91-3.28 | 0.86 | 0.60-1.23 |
| Low | 98 (24.8) | 141 (25.9) | 147 (26.0) | 207 (30.0) | 140 (26.8) | 1.17 | 0.99-1.39 | 0.93 | 0.73-1.17 |
| Normal | 98 (24.8) | 103 (18.9) | 127 (22.4) | 124 (18.0) | 87 (16.7) | 0.89 | 0.74-1.08 | 0.91 | 0.71-1.15 |
| High | 47 (11.9) | 70 (12.8) | 64 (11.3) | 85 (12.3) | 46 (8.8) | 0.94 | 0.75-1.18 | 0.93 | 0.71-1.21 |
| Very high | 127 (32.2) | 211 (38.7) | 192 (33.9) | 214 (31.0) | 165 (31.6) | 1 |  | 1 |  |
| Skin surface sebum U-zone |  |  |  |  |  |  |  |  |  |
| Very low | 111 (28.1) | 147 (27.0) | 148 (26.1) | 232 (33.6) | 240 (46.0) | 1.55 | 1.24-1.94 | 1.57 | 1.16-2.11 |
| Low | 169 (42.8) | 211 (38.7) | 242 (42.8) | 247 (35.8) | 141 (27.0) | 0.87 | 0.70-1.08 | 1.31 | 0.99-1.73 |
| Normal | 50 (12.7) | 72 (13.2) | 76 (13.4) | 82 (11.9) | 49 (9.4) | 0.92 | 0.71-1.20 | 1.13 | 0.82-1.54 |
| High | 15 (3.8) | 29 (5.3) | 33 (5.8) | 45 (6.5) | 26 (5.0) | 1.24 | 0.89-1.73 | 1.34 | 0.91-1.97 |
| Very high | 50 (12.7) | 86 (15.8) | 67 (11.8) | 84 (12.2) | 66 (12.6) | 1 |  | 1 |  |
| Skin surface moisture T-zone |  |  |  |  |  |  |  |  |  |
| Very low | 10 (2.5) | 21 (3.9) | 33 (5.8) | 33 (4.8) | 22 (4.2) | 1.42 | 1.03-1.97 | 2.10 | 1.42-3.11 |
| Low | 29 (7.4) | 61 (11.2) | 46 (8.1) | 86 (12.5) | 75 (14.4) | 1.67 | 1.32-2.10 | 1.91 | 1.45-2.54 |
| Medium | 179 (45.4) | 246 (45.2) | 260 (45.9) | 296 (42.9) | 271 (51.9) | 1.26 | 1.09-1.46 | 1.42 | 1.18-1.72 |
| Normal | 176 (44.7) | 216 (39.7) | 227 (40.1) | 275 (39.9) | 154 (29.5) | 1 |  | 1 |  |
| Skin surface moisture U-zone |  |  |  |  |  |  |  |  |  |
| Very low | 96 (24.4) | 148 (27.2) | 119 (21.0) | 155 (22.5) | 137 (26.2) | 1.38 | 1.10-1.72 | 1.08 | 0.80-1.45 |
| Low | 98 (24.9) | 142 (26.1) | 167 (29.5) | 215 (31.2) | 198 (37.9) | 1.80 | 1.45-2.23 | 1.25 | 0.95-1.65 |
| Medium | 138 (35.0) | 173 (31.7) | 186 (32.9) | 232 (33.6) | 148 (28.4) | 1.29 | 1.05-1.60 | 1.15 | 0.89-1.45 |
| Normal | 62 (15.7) | 82 (15.0) | 94 (16.6) | 88 (12.8) | 39 (7.5) | 1 |  | 1 |  |
| Pigmentation |  |  |  |  |  |  |  |  |  |
| Dark | 235 (59.7) | 266 (48.9) | 254 (44.8) | 284 (41.2) | 154 (29.5) | 0.546 | 0.45-0.67 | 0.97 | 0.78-1.22 |
| Medium | 109 (27.8) | 181 (33.3) | 230 (40.7) | 318 (46.2) | 243 (46.7) | 1.052 | 0.86-1.28 | 1.11 | 0.88-1.38 |
| Light | 49 (12.4) | 97 (17.8) | 82 (14.5) | 87 (12.6) | 124 (23.8) | 1 |  | 1 |  |
| Skin pore size |  |  |  |  |  |  |  |  |  |
| Very large | 227 (58.5) | 300 (55.8) | 363 (64.6) | 443 (64.6) | 359 (68.9) | 1.845 | 1.36-2.51 | 1.15 | 0.81-1.62 |
| Large | 53 (13.7) | 65 (12.1) | 72 (12.8) | 74 (10.8) | 51 (9.8) | 1.366 | 0.96-1.95 | 1.06 | 0.72-1.58 |
| Medium | 47 (12.1) | 74 (13.8) | 56 (10.0) | 81 (11.8) | 56 (10.7) | 1.487 | 1.04-2.12 | 1.22 | 0.82-1.81 |
| Small | 34 (8.8) | 60 (11.2) | 46 (8.2) | 59 (8.6) | 38 (7.3) | 1.387 | 0.96-2.01 | 1.02 | 0.68-1.54 |
| Normal | 27 (7.0) | 39 (7.2) | 25 (4.4) | 29 (4.2) | 17 (3.3) | 1 |  | 1 |  |
| Eye wrinkle |  |  |  |  |  |  |  |  |  |
| Very large | 58 (14.7) | 107 (19.6) | 191 (33.7) | 386 (55.9) | 379 (72.6) | 22.23 | 16.9-29.3 | 2.54 | 1.86-3.47 |
| Large | 22 (5.6) | 53 (9.7) | 72 (12.7) | 87 (12.6) | 47 (9.0) | 9.53 | 6.91-13.18 | 2.05 | 1.45-2.92 |
| Medium | 71 (18.0) | 123 (22.6) | 136 (24.0) | 137 (19.9) | 60 (11.5) | 6.07 | 4.56-8.10 | 1.84 | 1.35-2.52 |
| Small | 141 (35.7) | 179 (32.8) | 134 (23.7) | 71 (10.3) | 34 (6.5) | 2.62 | 1.98-3.47 | 1.31 | 0.96-1.77 |
| Very small | 103 (26.1) | 83 (15.3) | 33 (5.8) | 9 (1.3) | 2 (0.4) | 1 |  | 1 |  |
| Melatonin, ng/L | 7.4±8.8 | 7.4±8.6 | 6.9±6.5 | 7.4±9.6 | 5.4±4.6 | 0.979 | 0.95-1.01 | 0.96 | 0.93-0.99 |
| Epidermal growth factor, pg/ml | 85.2±96.2 | 89.6±95.5 | 87.6±78.9 | 97.4±112.1 | 67.2±48.7 | 1.001 | 0.99-1.00 | 1.004 | 1.00-1.01 |

\*-adjusted for age, gender, skin type and other skin physiological parameters using final ordinal
regression model.

Higher skin aging grades were significantly associated with very low skin surface moisture at T-zone (aOR =2.10, 95% CI 1.42-3.11, compared normal moisture level) and with low skin surface moisture at U-zone (aOR=1.25, 95% CI 0.95-1.65, compared with high moisture). On the other hand, having very low skin surface sebum at the T-zone was protective against skin aging. Conversely, having very low skin surface sebum at the U-zone increased the OR of having higher skin aging grades by more than 50% compared to normal skin surface sebum (aOR=1.57, 95% CI 1.16-2.11). Having large wide eye wrinkles raised the risk of skin aging by 2.5-fold compared to those with fine narrow wrinkles (aOR=2.54, CI 1.86-3.47). Skin pigmentation and pore size were not significantly associated with the skin aging grades. Serum melatonin and epidermal growth factor levels were identified for the 541 study subjects and hence could not be included in the overall multivariate model. The lowest mean value of serum melatonin was 5.4±4.6 ng/L found in participants with grade 6 skin aging compared to those with grade 2 changes. The mean level of serum EGF was 85.2±96.2 pg/ml for people with grade 2 changes, in contrast to 67.2±48.7 pg/ml for those with grade 6 skin changes. However, on the crude analysis, decreasing serum melatonin levels by one unit was associated with an approximately 4% increase in the odds of increasing their skin aging grade (OR=0.96, CI 0.93-0.99).

Figure 2 includes correlations found between the skin physiological parameters, age, body mass index, sun exposure time, serum melatonin and EGF and skin aging grades. We used spearman’s correlation analysis because of our ordinal dependent and skew distributed continuous independent variables. The skin aging grade was strongly positively correlated with age (rs=0.801, p<0.01). There was a weakly positive correlation between body mass index and skin aging grades (rs=0.26, p<0.01). Surprisingly, the relationship between skin aging grade and sun exposure time was negligible. The width of eye wrinkles was moderately directly correlated with both skin aging grade and age (rs=0.50, p<0.01 and rs=0.54, p<0.01, respectively). There was a weak but positive correlation between skin pore size and skin surface sebum at T-zone (rs=0.21, p<0.01), whereas a negative correlation between skin pore size and surface moisture at T-zone was identified (rs=-0.075, p<0.01).

**Fig 2.**
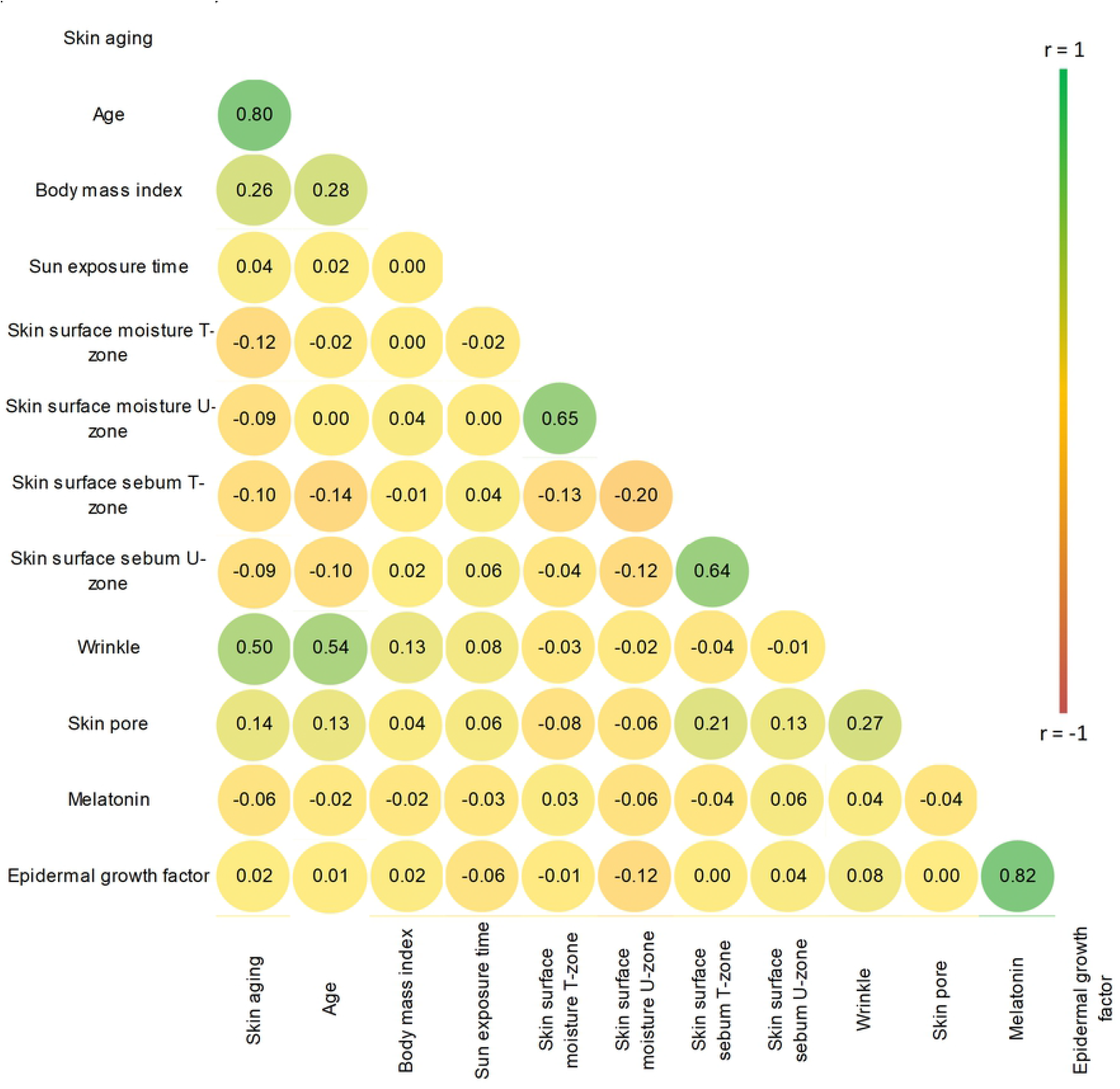
Correlation between skin physiological measurements and skin aging grades for study participants (N=541).

## Discussion

Our results indicate that being older, living in an urban area, having a lower frequency of getting professional skin care services, not using sunscreen, working outside, working in harsh conditions, smoking, having a higher BMI, and more sun exposure were significantly associated with higher skin aging grades. According to our skin physiological parameters, being very low surface moisture at T- and U-zones and having a dry skin type were significantly related to skin aging.

According to Green et al., every extra year of age after 30 up to 54 years significantly increased the odds of having a higher skin aging grade by 14%, and people frequently working outdoors had 60% higher odds [7]. These findings are similar to our study and other previous published study [12, 20, 21]. Research in Australia illustrated that the odds of skin aging in men were threefold higher than for women [20]. In our study, gender had no significant effect on skin aging grade, and this is probably due to Mongolia’s long cold winters during which there is comparatively little sunlight. In a Queensland population 20-55 years of age, outdoor occupations and outdoor leisure activities were crudely associated with higher skin aging grades [22]. This is similar to our study results where we found working outside was associated with a 1.4-fold increase in skin aging compared to those working in optimal conditions. Also, working in harsh conditions in Mongolia had a similar effect on those working in the same conditions in Australia.

Among identical twins, smoking has been identified as an independent risk factor in skin aging, concluding that a 5-year difference in smoking duration is associated with skin changes [21]. In a study conducted among the Brazilian population to assess the relationship between tobacco use and skin aging, a smoker using 40 packs per year had 3.9 times higher skin aging grades than non-smokers [23]. However, our smoking data were not granular and did not include the number of pack-years of cigarettes in current and former smokers. Nonetheless, being a current smoker increased the odds of skin aging by 30 percent. Several studies concluded that people with high BMI were significantly more likely to have higher skin aging grades on crude analysis, but not after adjusting for age and sex [7, 20], consistent with our study. A clinical study of postmenopausal women concluded that women who slept for 5 hours per day had higher trans-epidermal water loss levels and decreased skin barrier recovery after UV-induced erythema [8]. Our study participants who slept 4-5 hours per day had higher odds of skin aging on crude analysis. According to a previous study, there was a moderately positive correlation between skin aging grade and sun exposure time [24]. In our findings, these variables were weakly correlated. Menopause-related skin changes lead to facial wrinkles and skin aging [25]. We found that postmenopausal women were 18 times more likely to have higher skin aging grades on crude analysis. But on multivariable analysis, this was not significant.

Besides determining potential personal, socioeconomic and environmental predictors resulting in skin aging, we assessed skin physiological parameters, including skin surface moisture and sebum, skin pigmentation and wrinkles and pore size. We found after adjusting for physiologic factors that people with very low moisture levels at the T- and U-zones were significantly more likely to have higher skin aging grades. Generally, those with the dry skin type had the highest odds of having higher grades than the normal skin type. A review concluded that one of the characteristics of extrinsic skin aging is dry skin, and having dry skin was associated with high skin aging grades [26]. According to a study involving southeast China residents, skin aging related to skin type, and people with oily skin had lower odds of facial aging compared to those with dry skin [13]. Moreover, melatonin and epidermal growth factor protect against DNA damage caused by UV radiation and prolongs the process of photoaging [19, 27]. We found that lower serum melatonin level was one of the risk factors leading to aging skin.

The method of evaluating skin aging employed in our study is appropriate for a population-based study because of its efficiency, low cost and convenience. We believe this is the first nationwide study to assess the possible factors that have been shown to affect the rate of development of skin aging in Mongolia. Most previous studies have been completed in an unrepresentative sample from clinical settings; our results are based on a representative sample of urban and rural communities instead of from those seeking care for dermatologic conditions.

Our study identified that where one lived in Mongolia affected skin aging and, surprisingly, people living in urban areas had higher odds of skin aging. Because living in Ulaanbaatar exposes people to Mongolia’s highest concentrations of particular matter derived from burning coal during the long winter season, from automobiles and coal-burning power plants year-round [28, 29], we hypothesize that air pollution ages the skin. A previous study investigating the association between airborne particles and skin aging among Caucasian women revealed that airborne exposure was strongly related to aging skin [30]. Therefore, we propose additional studies to clarify further the association between air pollution and skin aging in Ulaanbaatar.

## Conclusions

Our study evaluated skin aging grades in a nationwide survey of Mongolians 18 years of age and older to determine skin-aging risk factors. Being older, living in an urban area, tobacco smoke exposure, having a dry skin type, more sun exposure, working in harsh conditions and working outside increase the odds of skin aging. Having very low skin surface moisture in the T- and U-zones was significantly related to higher odds of skin aging. Our findings provide a rationale to design preventive strategies for reducing skin-aging changes in young and middle-aged adults’ skin. The principal preventative methods are avoidance of sun exposure without protection, avoiding tobacco smoke.

## Acknowledgments

We extend our sincere gratitude to the Health Care Centers of Uvs, Khuvsgul, Umnugovi and Dornod provinces for their practical support throughout the study. We thank all the study participants for participating and sharing their insights that made the study’s success possible. We thank Ron Anderson MD for the critically reviewing and editing our manuscript. We thank Nasantogtox Erdenebileg, MD for assisting statistical analysis.

## Supporting information

### S1 Appendix

Questionnaire in both English and Mongolian language.

